# RiboDiPA: A Novel tool for differential pattern analysis in Ribo-seq data

**DOI:** 10.1101/2020.04.20.050559

**Authors:** Keren Li, Matthew Hope, Xiaozhong A. Wang, Ji-Ping Wang

## Abstract

Ribosome profiling (also known as Ribo-seq) has become an important technique to investigate changes in translation across a wide variety of contexts. Ribo-seq data not only provides the abundance of ribosomes bound to transcripts, but also positional information across transcripts that could be indicative of differences in translation dynamics between conditions. While many computational tools exist for the analysis of Ribo-seq data, including those that assess differences in translational efficiency between conditions, no tool currently exists for rigorous test of the pattern differences in ribosome footprint. In this paper we propose a novel approach together with an R package, RiboDiPA, for <u>D</u>ifferential <u>P</u>Pattern <u>A</u>nalysis of Ribo-seq data. RiboDiPA allows for quick identification of genes with statistically significant differences in ribosome occupancy patterns for model organisms ranging from yeast to mammals. We show that differential pattern analysis reveals information that is distinct and complimentary to the existing methods that focus on translational efficiency analysis. Using both simulated Ribo-seq footprint data and two benchmark data sets, we illustrate that RiboDiPA can not only uncover meaningful global translational differences between conditions, but also the detailed differential ribosome binding patterns to a single codon resolution.

## 1 Introduction

Translation of mRNA messages into proteins is a fundamental process that all organisms must undergo in order to survive. Given its importance, translation is highly regulated by organisms across all kingdoms of life to ensure proper protein expression in the right contexts [1], while mis-translation or mis-regulation of translation has frequently been linked to disease [2]. Protein production itself is carried out by ribosomes as well as key accessory protein complexes, which are conserved across species. Molecular and genetic dissection of translation has revealed the details of a complex process consisting of multiple steps, from initiation to elongation to termination and recycling. Translation initiation is perhaps the most-well characterized of these steps [3], although regulation at later steps has also been demonstrated to be critical [4]. Importantly, cells have evolved strategies to adaptively regulate translation of mRNA messages in changing environments, including under various types of cellular stress, such as oxidative stress and starvation [5].

In the past decade, a new method called ribosome profiling (or Ribo-seq) has harnessed the power of next-generation sequencing technologies to provide a new way of quantitatively examining translation on a gene-by-gene basis [6]. Ribosome profiling not only offers quantification of translational efficiency of a gene, which is the number of ribosomes per mRNA molecule under a given condition, but also important information about the distribution of ribosomes across an mRNA transcript. This shape information has been used to identify upstream open reading frames (uORFs) [7], observe stop codon read-through [8], and determine the sites of pausing during translation [9], among many other applications. Ribosome profiling has been applied to many organisms [10] and improvements upon the original technique have allowed for more precise targeting of translational events [11].

As the popularity of ribosome profiling as an investigative technique has increased, the computational tools and approaches to analyze Ribo-seq data have rapidly proliferated. The purposes of these tools range from data preprocessing, uORF discovery and annotation, quantification of translational efficiency, etc. For extensive and excellent reviews of these tools, we refer to [11, 12, 13]. In this paper, we are particularly interested in statistical inference of translational differences between conditions using Ribo-seq data. In this regard, there are several tools available for differentiation of translational efficiency between conditions based on the gene-wise Ribo-seq counts [14, 15, 16, 17, 18, 19, 20, 21]. The translational efficiency for each gene is typically quantified by Ribo-seq read count normalized by the RNA-seq gene expression. While these approaches can provide insight into translational efficiency, the ribosome binding pattern along the gene body, which we believe can provide a more complete picture of transcriptional and translational variations across conditions, has been mostly ignored. For example, a given gene with similar translational efficiency quantified by total Ribo-seq and RNA-seq counts may have differential Ribo-seq read distribution pattern along the transcript, which may be indicative of difference in translational mechanisms of interest. Nevertheless there is yet no existing method or tool for rigorous test of the differences of ribosome binding patterns. In this paper, we propose a novel method together with an R package named RiboDiPA for differential pattern analysis in Ribo-seq data. We will show this approach can uncover new mechanisms of translational regulation in Ribo-seq footprint data. The new approach will be illustrated using a systematic simulation study and two benchmark data sets.

## 2 MATERIALS AND METHODS

### 2.1 Experimental data sets

We selected two experimental case studies to assess how our method performs. The first one is a high quality dataset collected in yeast by [22]. This study used multiple translation inhibitors to reveal changes in ribosome occupancy upon various cell stresses, and defined classes of RPFs with different lengths corresponding to ribosomes with open or occupied A-sites. In this study, we shall compare unstressed yeast to yeast challenged with three different stresses– osmotic stress, oxidative stress, and stationary phase growth– as well as compare wild type yeast to yeast of different genetic backgrounds, ΔeRF1 and ΔRck2. The second data set comes from a mouse study [23] which compares mouse embryonic stem cells with different dosages of the translational regulator Nat1: wild-type, heterozygous mutant, and null. These data sets were selected because they represent two divergent model organisms of typical sample size (2-3 replicates per condition), and more importantly, both have relatively high read counts.

### 2.2 Overview of RiboDiPA flow

RiboDiPA, implemented in an R package, provides a pipeline for pattern differentiation for Ribo-seq data. Figure 1 provides a flow chart of RiboDiPA pipeline from exon concatenation, BAM file processing, P-site mapping, to differential pattern test. The package takes the Ribo-seq alignment file (in .bam format) and Genome Transfer File (.GTF) as inputs and outputs the differential pattern (DP) analysis results with statistical significance and supplementary pattern dissimilarity measure (*T*-value). In addition, it provides additional functions for visualization of Ribo-seq footprints with specified resolution. In terms of computing time, it takes about 10 minutes to run the entire pipeline for the wild type vs. eRF1d comparison of yeast data (4 samples) at single-codon resolution on a 20-core node on a Linux cluster. RiboDiPA can be downloaded at github (https://github.com/jipingw/RiboDiPA). The details of each function of the package are described below.

**Figure 1:**
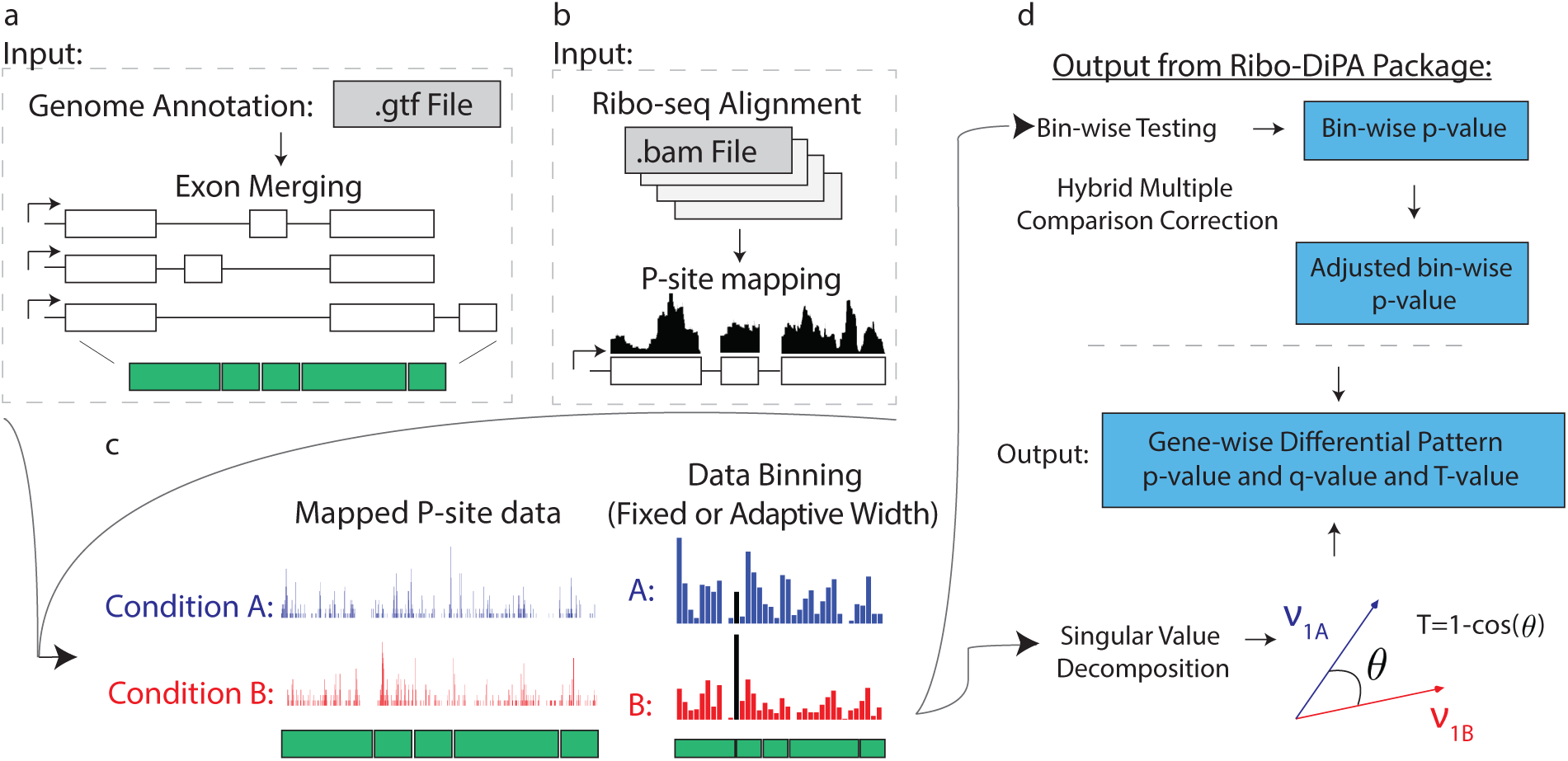
RiboDiPA package workflow. Input for the RiboDiPA package are: (a) the Genome Transfer File (GTF) of the experimental organism and (b) Ribo-seq alignment files in BAM format with one file per replicate. All exons, 5’UTR(s), and 3’UTR(s) from the same gene are concatenated to form a total transcript. RPFs are parsed and the P-site position is calculated for each RPF. (c) Mapped P-site data representing the P-site frequency at each nucleotide position along the total transcript (left) and the binned P-site data with customizable bin width (right). (d) Flow of differential pattern analysis and output of RiboDiPA including *p*-value, *q*-value and *T*-value for each gene under testing.

### 2.3 Exon concatenation, P-site mapping and data binning

Since true RPFs originate from ribosomes that are actively translating in coding regions, in cases where a gene has multiple splicing isoforms, it is difficult to distinguish which isoform an observed Ribo-seq read comes from. Our pattern analysis is performed on the gene-level by concatenating all exons from the same gene into a total transcript in order to get a merged picture of translation. We first mapped all RPFs to the genome, and then identified the P-site position of each RPF in the corresponding total transcript (see Supplementary Materials). The aggregated RPF count at each codon was binned at specified bin width for the downstream pattern analyses.

RiboDiPA allows for differential pattern analysis of Ribo-seq data with customizable bin width. The motivation for data binning is that the read coverage in Ribo-seq experiments tends to be sparse at invididual codons. The differential pattern analysis to be proposed below requires testing differential mean of read counts at each location of the transcript. Binning helps alleviate excessive statistical tests on locations with 0 or extremely low counts. More importantly, more position-wise tests within the same gene increases the chance of type I error, and correction for which may undermine the power (to be further discussed below). In RiboDiPA the user can specify a fixed bin width as small as a single codon (3 nt). In addition we have implemented an adaptive bin width selection approach by Freedman and Diaconis [24] originally proposed for histogram construction. Our simulation results suggest that Freedman-Diaconis rule empirically outperforms several other popular approaches including Sturges’ formula [25], Doane’s formula [26], Rice Rule [27], Scott’s normal reference rule [28] and an improved Doane’s formula using a kurtosis criterion [29, 30) (see Simulation section]. Briefly, suppose there are *m* P-sites mapped at locations *x*_1_, …, *x*_*m*_ of a transcript. The adaptive bin width is calculated as follows:

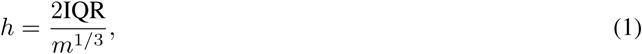

where IQR is the interquartile range of the data. In RiboDiPA, we first calculate the average P-site frequency at each position of the transcript across replicates. The adaptive bin width is calculated based on the average footprint using formula (1). The common empty bins shared by all replicates are removed after binning in RiboDiPA. In the following context, we shall refer to bin interchangeably as “position” or “location”.

Lastly, we only consider genes with at least one Ribo-seq read in every replicate for illustration of differential pattern analysis throughout this paper.

### 2.4 Differential pattern analysis

We begin with two examples from a yeast translation study [22] to demonstrate the necessity to develop new methods for differential pattern analysis in Ribo-seq data. A naive test for differential translation is to test for mean or abundance equivalence based on the gene-wise total Ribo-seq read counts (equivalent to differential expression analysis in the RNA-seq data). To avoid confusion below, we shall refer to this test as differential abundance (DA) analysis. Figure 2 shows the ribosome footprint of two genes, plotted in the form of mapped P-site frequency at each nucleotide location (unbinned) or at each bin (adaptively binned) along the transcript, for unstressed yeast cells and cells undergoing oxidative stress, in blue and red respectively with two replicates per condition. Neither TCB3 (YML072C) nor ABC1 (YGR037C) gene showed significant difference in terms of overall RPF abundance between conditions (*p*-value = 0.995 and 0.669 respectively). In contrast, TCB3 shows similar ribosome occupancy patterns between unstressed and stressed cells while ABC1 shows a dramatic shift in the distribution of ribosome occupancy toward the middle of the gene under oxidative stress. Clearly, different ribosome binding pattern cannot be necessarily inferred by differential abundance analysis using total read counts, therefore we set out to establish a new framework for testing pattern differences in ribosome profiling data.

**Figure 2:**
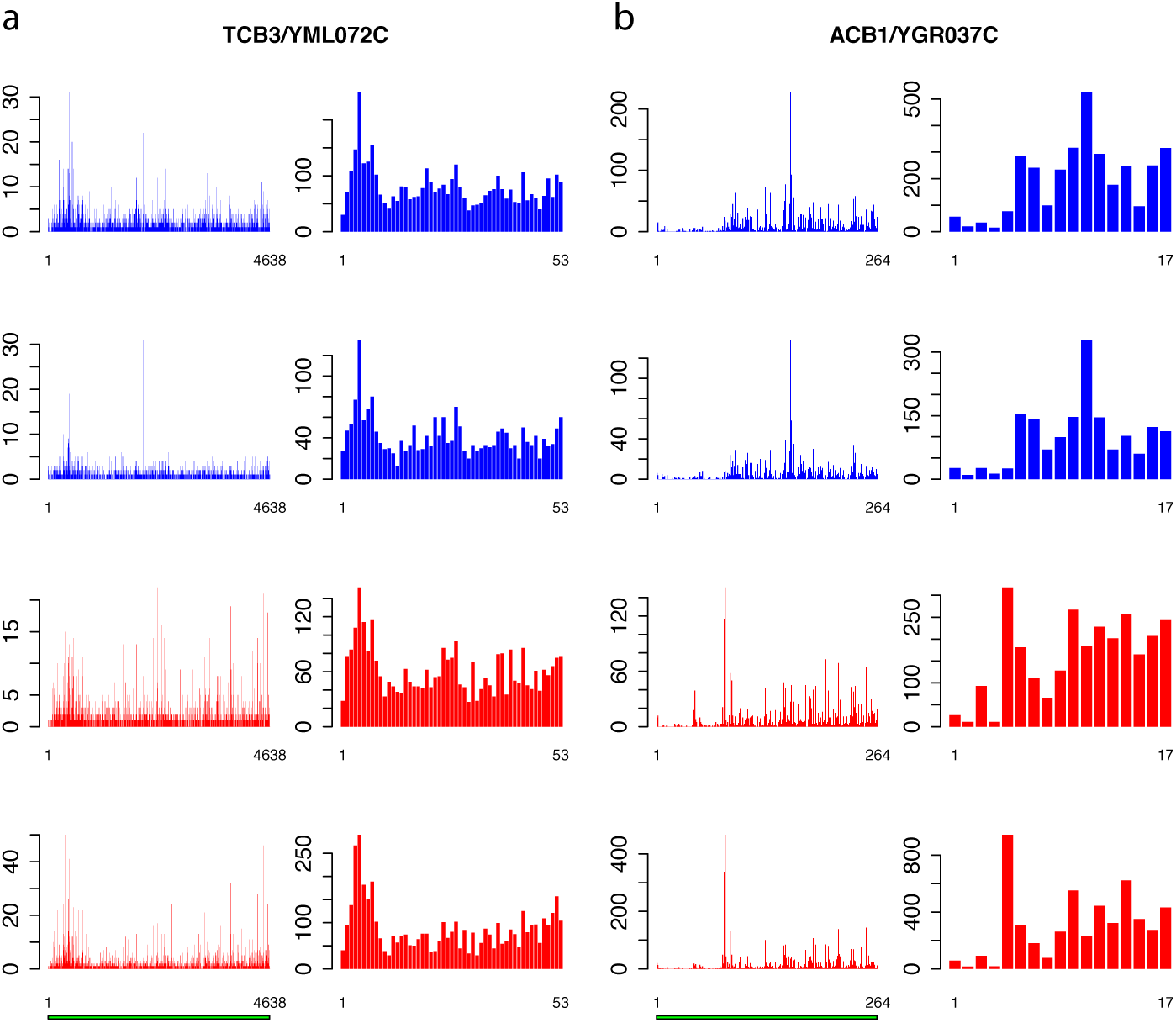
Ribosome profiling data shows example genes with differences in ribosome occupancy patterns, but similar abundance of RPFs. Plotted are the ribosome profiling data for two example genes from yeast [22], TCB3 (a) andACB1 (b) with two replicates in unstressed conditions (blue) and under oxidative stress (red). Both TCB3 and ACB1 do not show a significant difference in the abundance of RPFs between conditions (DA test defined in text), but ACB1 shows a clear pattern difference of ribosome occupancy near 5’ end whereas TCB3 does not. For each panel, the distribution of ribosome unbinned P-site frequency is shown on the left and the binned data on the right.

To define differential pattern (DP) analysis rigorously, we propose a statistical framework as follows. Suppose we have two conditions with *n*_1_ and *n*_2_ replicates respectively. For a given gene *g* of length *J*, denote the read count at position *j* in the *i*th replicate of condition *k* as *X*_*ijk*_, for *j* = 1, …, *J, i* = 1, …, *n*_*k*_, *k* = 1, 2 (we omit the gene index in notations for simplicity). We assume

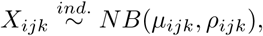

for *i* = 1, …, *n*_*k*_, *j* = 1, …, *J, k* = 1, 2, where

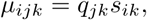

and *ρ*_*ijk*_ is the dispersion parameter. Note that *s*_*ik*_ is a gene- and sample-specific scale factor measuring the RPF abundance in the *i*th replicate in the *k*th condition, and *q*_*jk*_ is a position- and condition-specific relative abundance parameter measuring ribosome binding affinity. For a given gene, the scale factor *s*_*ik*_ can vary across replicates and conditions (and hence mean *µ*_*ijk*_) due to differences in abundance level and sequencing depth. Our differential pattern analysis for a given gene is defined for testing the following hypothesis:

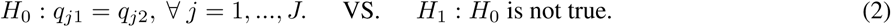

In words, the pattern equivalence can be interpreted as having parallel mean curves (*µ*_*ijk*_ defined for *j* = 1, .., *J*) across different conditions.

The null hypothesis specified in (2) can be regarded as intersection of a set of sub-hypotheses, i.e., 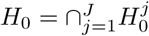 where 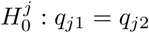, for *j* = 1, …, *J*. This leads us to consider a point-wise approach by first testing each sub-hypothesis separately, i.e.,

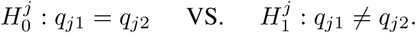

We convert this test into a standard differential expression analysis problem by treating *s*_*ik*_ as a sample- and gene-specific normalizing factor. Thus the first step is to estimate the normalization factor *s*_*ik*_ for each replicate of each gene. In RNA-seq analysis, there are many existing methods for calculation of normalization constants. For example, Anders and Huber [31] and Love et al [32] took the median of the ratios of counts to geometric mean of counts across samples as a normalizing constant to avoid the influence of highly and differentially expressed genes:

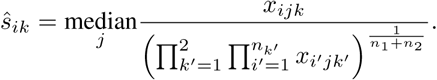

For the Ribo-seq data, the read count at each location for a given gene can still be very low even after binning, often causing near 0 normalization constants *ŝ*_*ik*_. In contrast, we found the total read count is a more robust measure of the abundance level. To exclude the bins that represent the true differential pattern, we defined an outlier bin as that whose log_2_-fold change value is more than 1.5 interquartile ranges (IQRs) below the first quartile or above the third quartile. Denote the remaining non-outlier bin set as *J*′. The normalizing constant is defined based on the total read counts from each replicate for the same gene, i.e.,

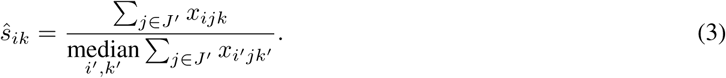

To perform a negative binomial test, one critical step is to estimate the dispersion parameter, typically modeled as a function of the mean. One challenge arises due to the small sample size such that the dispersion parameter cannot be well estimated on a gene-by-gene basis. Instead the gene specific dispersion parameter is estimated by aggregating information from all genes. For instance, Robinson and Smyth [33] first estimated the gene-wise dispersion parameter from the conditional maximum likelihood conditioning on the total count for that gene, and then shrunk it towards a consensus value by an empirical Bayes model (“edgeR” R package). Anders and Huber [31] assumed a locally linear relationship between dispersion and mean to borrow information across genes (‘“DESeq” R package). As an update, Love et al [32] first used gene-wise maximum likelihood estimate (MLE) of the dispersion parameter to fit a dispersion-mean curve, the fitted value of which was further used as the mean of a prior distribution to produce an estimate of gene-specific dispersion parameter shrunk towards the mean curve (“DESeq2”). In this paper we adopt the method in DEseq2 to estimate the bin-wise dispersion parameter for its established competitive performance. For the same gene, the relatively small bin number (median is 14 for the yeast data) makes it impossible to robustly estimate of gene-specific dispersion-mean function. Hence we pool all bins from all genes and feed the read count *x*_*ijk*_ together with corresponding scale normalization factor *ŝ*_*ik*_ into DESeq2 for estimation of the common dispersion-mean function, and thereafter the dispersion parameter for each bin within each gene. The bin-wise test for differential mean was subsequently carried out using DESeq2.

Let *p*_*j*_ be the DESeq2 *p*-value for the *j*th bin of a given gene for *j* = 1, …, *J*. The type I error for testing original hypotheses *H*_0_ VS. *H*_1_ defined in (2) equals Prob(reject at least one 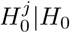), i.e., the family-wise error rate for testing 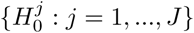. Thus to control the type I error rate for testing *H*_0_ VS. *H*_1_, we need to control family-wise type I error rate in the point-wise approach.

The well-known but conservative Bonferroni procedure is to adjust every *p*-value *p*_*j*_ to min(*Jp*_*j*_, 1). Here we follow a recently proposed hybrid Hochberg-Hommel method by Gou and coauthors [34] (implemented in R package “elitism”) to calculate the adjusted *p*-values for each sub-hypothesis for its relatively more powerful performance. Briefly, let *p*_[1]_ ≥ … ≥ *p*_[*J*]_ be the ordered *p*-values. The adjusted bin-level *p*-values is given by

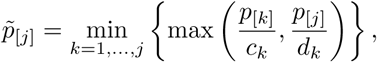

with *c*_*k*_ = (*k* + 1)*/*(2*k*) and *d*_*k*_ = 1*/k*. The gene-level *p*-value is defined as 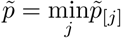.

Denote 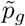 as the gene-level *p*-value for gene *g*, for *g* = 1, …, *G*. For false discovery rate control, we input the gene-level *p*-value into *q*-value package [35] to calculate the *q*-value for each gene.

### 2.5 A supplementary pattern dissimilarity measure by SVD

The gene level *p*-value from the hybrid Hochberg-Hommel procedure only reflects the statistical significance at one individual location that achieves the the minimum *p*-value among *J* locations/bins. It is less indicative about the read count scale at the differential bin(s) or how many bins have differential patterns. To provide a supplementary and more direct measure of pattern dissimilarity, we consider an alternative approach based on singular value decomposition.

Let 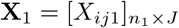 and 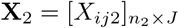 be the count matrices from condition 1 and condition 2 respectively where each row stands for one sample and each column for one position. It is well known that the first right singular vector from singular value decomposition (SVD) of a matrix represents the first principal row pattern. Thus if **X**_1_ and **X**_2_ preserve the same pattern, we expect to see similar first right singular vectors. Let **v**_1_ and 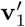 be the first right singular vectors from sample **X**_1_ and **X**_2_ respectively. Define the *T* value as follows:

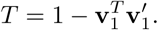

Note that 0 ≤ *T* ≤ 1. Geometrically, it equals the 1-*cosine* of the angle of **v**_1_ and 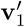 (Figure 1d). A larger *T* may provide more evidence to reject *H*_0_. There are some known results on the null distribution of *T* when *X*_*ijk*_ are i.i.d. Gaussian, which however are not applicable to non-i.i.d. count data here. In our case, *T* is not a pivotal quantity under *H*_0_ and its null distribution depends on the actual values of mean and variance at each location. In the RiboDiPA package, we output *T* value as a complementary statistic for users to prioritize positive genes with differential patterns (analogous to fold change in differential expression analysis). We shall demonstrate its usefulness below from both real and simulated data.

## 3 RESULTS AND DISCUSSION

### 3.1 Differential pattern (DP) vs. differential abundance (DA) analysis in Ribo-seq

To provide an overview of how DP analysis differs from DA analysis, we performed both analyses on WT unstressed yeast cells versus WT cells experiencing oxidative stress from the yeast data. Both DP and non-DP groups showed similar patterns of scatter plots of read counts (Figure 3a,b), though genes identified with DP tended to have larger average read counts. This is not surprising as the DP test is based on bin-wise tests and fewer reads within a bin may result in inadequate power for detection of differences due to large dispersion. We discovered that many genes showed differential patterns in ribosome occupancy, while few of which showed differential abundance (Figure 3a,c). For example, out of the 5,746 genes that met our criteria for analysis, 478 were significant in DP but not in DA; and only 18 were significant in both DA and DP analyses (q-value ≤ 0.05). These results suggest that DA analysis cannot be used for pattern differentiation in the Ribo-seq data.

**Figure 3:**
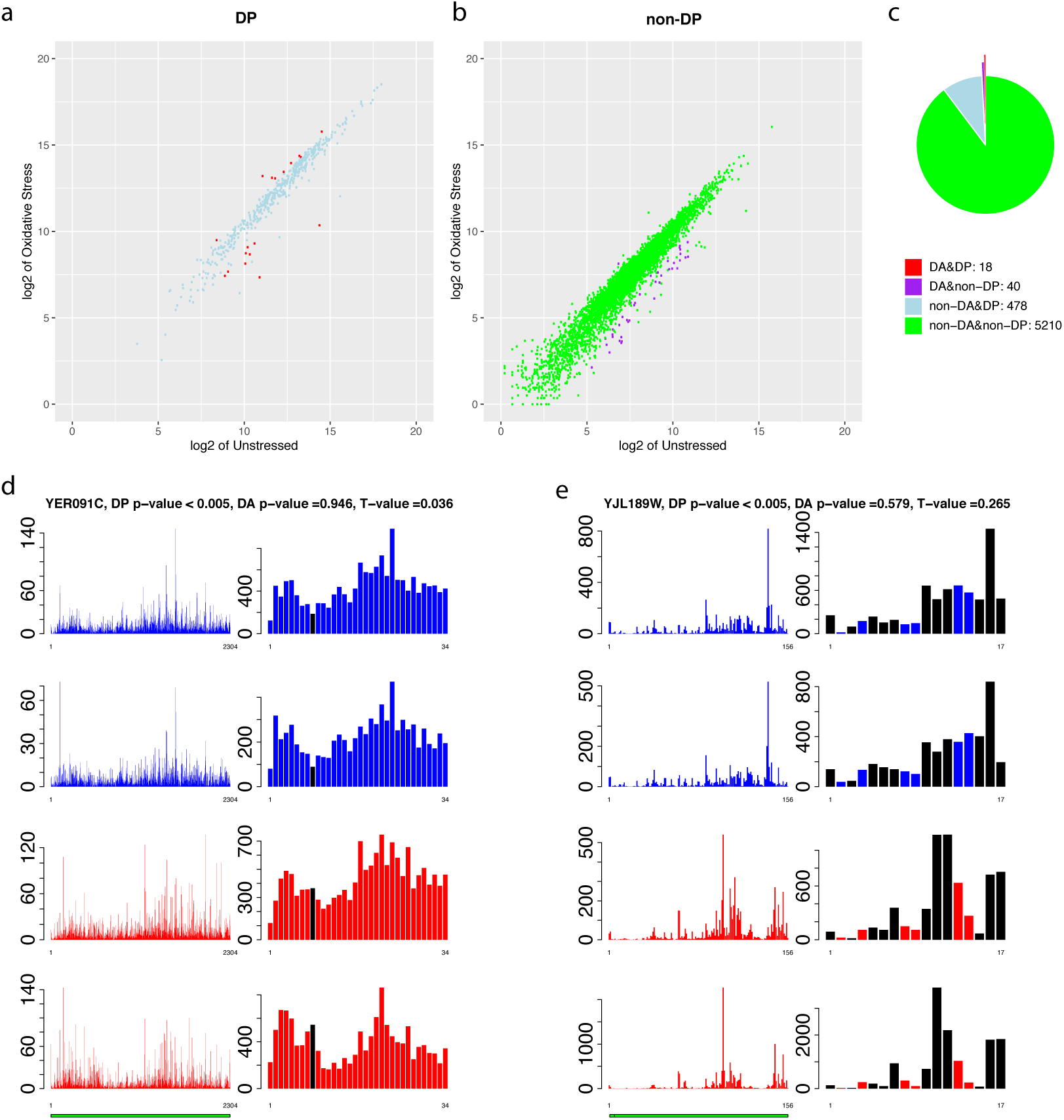
Differential pattern (DP) vs. differential abundance (DA) analysis. Data throughout the figure concerns WT unstressed yeast cells and WT cells responding to oxidative stress from yeast data set. (a) Scatter plots of average Ribo-seq read counts for genes with DP and (b) without DP (at q-value ≤ 0.05). (c) Pie chart for genes tested significant/insignificant in DP/DA analysis. Among 5,746 genes analyzed, 5,250 genes had no differential pattern (of which 40 were DA negative), however 496 genes had a significant differential pattern (of which 478 were DA negative). Panels (d) and (e) present the footprints of two genes(both unbinned and binned, unstressed in blue, stressed in red) to show that *T*-value can be used as a supplementary measure to identify genes with larger pattern differences beyond statistical significance measure *p*-value or *q*-value. Bins colored in black are those having significant adjusted *p*-value ≤ 0.05 in the DP test.

### 3.2 *T*-value as a supplementary measure for pattern difference

RiboDiPA outputs a list of significant genes with pattern difference. Practically, we are more interested to identify genes with DP in more bins/positions or in bin(s) with larger read counts. The gene-wise *p*-value is not informative in this regard as it is dictated by the most significant bin/position in the gene and it does not tell the read count magnitude or how many bins are DP significant. Analogous to log fold change used in gene expression analysis, the *T*-value provides a metric to prioritize investigation of genes that have more pronounced changes in ribosomal occupancy patterns. We show two example genes from our oxidiative stress analysis that tested positive for DP between the two conditions: MET6 (YER091C), a gene involved in methionine metabolism, and RPL39 (YJL189W), a large-subunit ribosomal protein. Figure 3d-e show the P-site footprints before (left) or after (right) binning, with WT unstressed replicates in blue and WT oxidative stress in red. While adjusted *p*-values of both genes are significant (<0.005), the *T*-value of MET6 is much smaller (0.036) than RPL39 (0.265). Examining the Ribo-seq footprints of RPL39 shows a dramatic shift in occupancy away from the 3′end of the gene towards the interior of the gene body. Therefore, ranking genes that test positive for DP by their *T*-value could be a valuable way for investigators to focus their attention on genes with the most pronounced changes between experimental conditions.

### 3.3 DP analysis uncovers global translational difference

To investigate how DP analysis can help gain biological insights into global translational differences between conditions, we examined four additional comparisons for the yeast data including: WT osmotic stress and WT starvation response (the latter known as stationary phase growth) vs. WT unstressed cells; and a Rck2 deletion strain vs. WT cells in both oxidative and osmotic stress. Rck2 is a critical kinase involved in stress response, which has been demonstrated by the authors to have effects on translation in the case of osmotic but not oxidative stress. We plotted the empirical cumulative distribution function (ECDF) of *p*-value for all five comparisons for the DP test and the respective number of test-significant genes as a function of *q*-value threshold (Figure 4a,b). Consistent with the previous authors’ findings, we found no significant genes with DP in the WT versus ΔRck2 comparison for oxidative stress at *q*-value threshold =0.05, while for the rest, substantial number of genes with differential occupancy patterns were identified. For example, at the same *q*-value threshold the method discovers hundreds of genes for the three stress conditions compared to unstressed cells, and approximately 110 genes for the WT to ΔRck2 comparison under osmotic stress (Figure 4b). These results provide global pictures of translational difference between different biological conditions.

**Figure 4:**
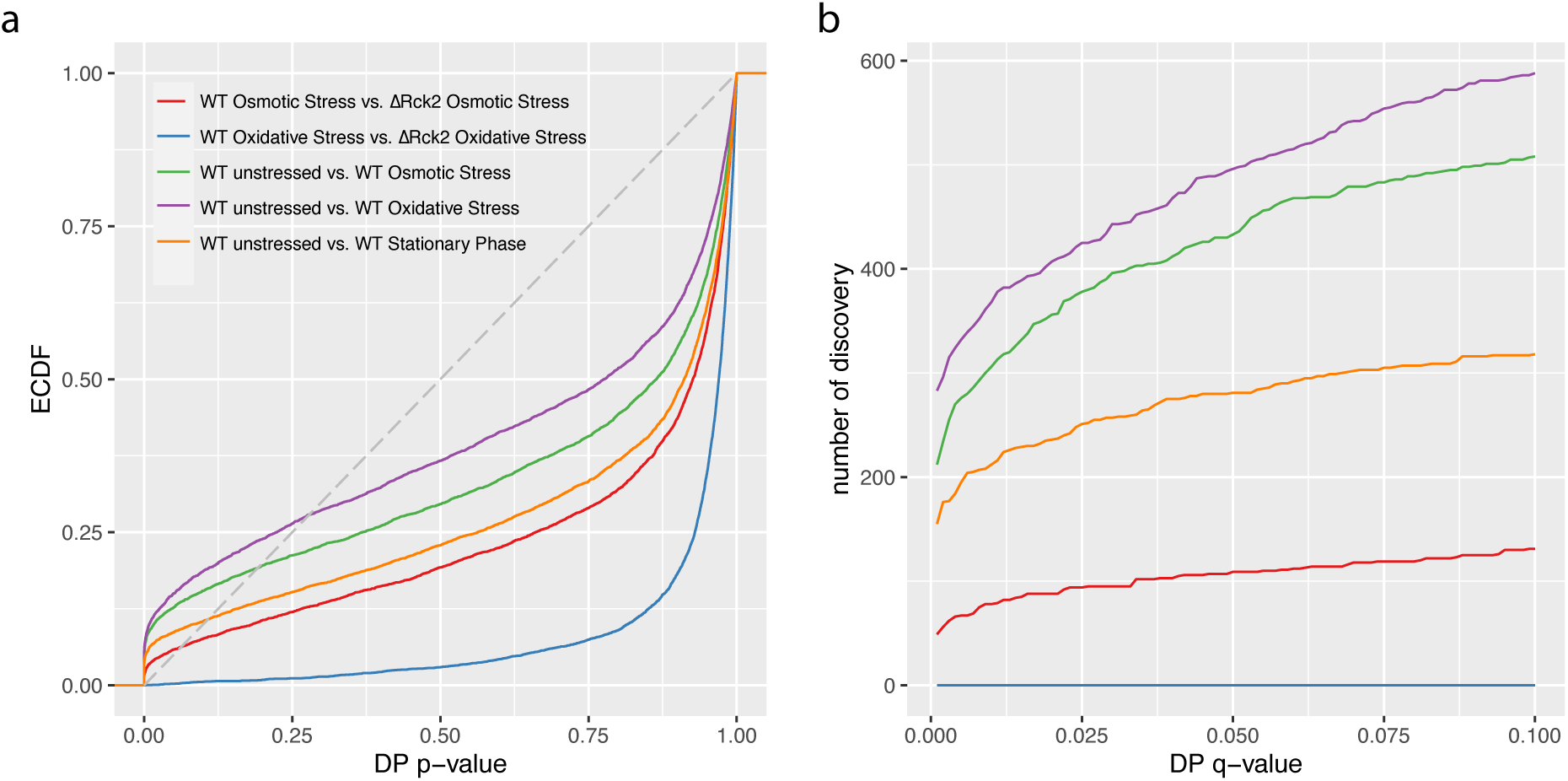
DP analysis shows global difference in translational activities between conditions. Plotted are (a) the empirical cumulative distribution function (ECDF) of *p*-value for DP analysis for five different comparisons of yeast stress conditions from the yeast data set, and (b) the corresponding number of discoveries under different *q*-value threshold values for each comparison.

### 3.4 RiboDiPA for single-codon resolution DP analysis

RiboDiPA allows to perform DP analysis at single-codon resolution to identify fine pattern differences between conditions. To illustrate this, we turned to a different data set from [22], which compares WT yeast cells (unstressed) to a Eukaryotic Release Factor 1 (eRF1) deletion strain (unstressed). eRF1 is a key factor in translation termination from all three stop codons in yeast, and the authors previously showed that eRF1 deletion causes accumulation in ribosome occupancy at two locations near the stop codons. Consistent with the conclusions of the previous authors, many genes showed significant pattern differences towards the 3’ end of the gene from DP analysis with adaptive binning (see Figure S.1 in the Supplementary Materials). To pinpoint the exact location of differential patterns, we carried out a single-codon DP analysis and examined the 50 codons down-/up-stream of the start and stop codons respectively. Figure 5a plots the number of genes that have significant DP at each given codon (adjusted condon-level *p*-value ≤ 0.05), where positive direction indicates enrichment of Ribo-seq reads in eRF1d relative to WT, and the negative for down-regulated occupancy in eRF1d. RiboDiPA precisely identified two large spikes at the second and twelfth codon upstream the stop codon, with all significant genes at these positions showing an increase in occupancy in the eRF1d condition relative to wild type. The fine details of the differential patterns were further exemplified in two genes, SUI3 and TEF2 in Figure 5b-c.

**Figure 5:**
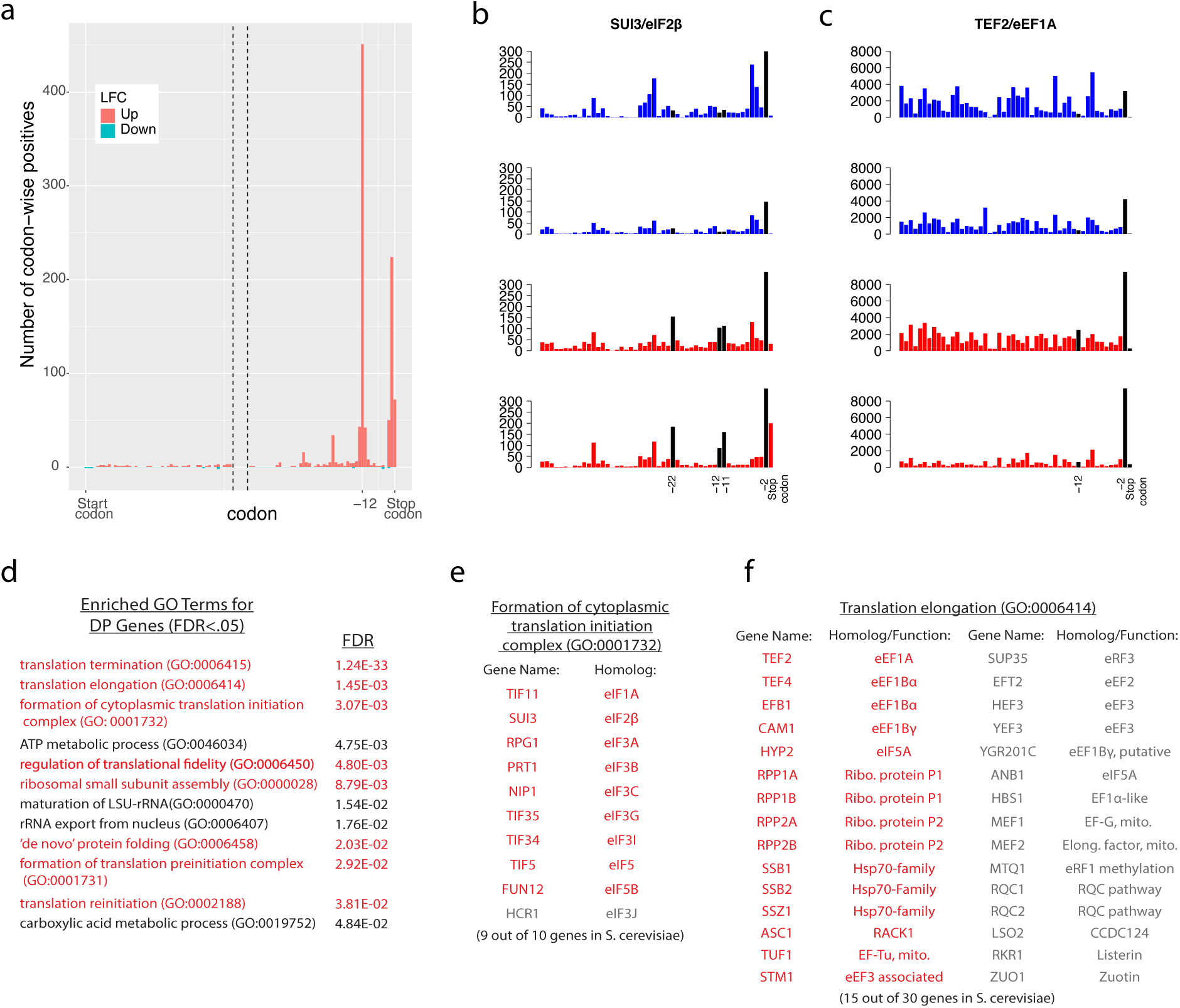
RiboDiPA for single-codon resolution DP analysis. (a) Comparison of WT unstressed yeast cells vs. eRF1 deletion strain (unstressed) from [22] shows significant enrichment of P-sites at the −12 and −2 codon position in the eRF1d cells (stop codon defined as −1 position), while no significant differential pattern is present around start codon. Plotted in vertical axis is the number of genes that have adjusted *p*-value ≤ 0.05 at each given codon, with positive direction for enrichment in eRF1d, and negative direction for depletion. Panels (b) and (c) show two example genes, SUI3, an eIF2*β* homolog, and TEF2, an eEF1A homolog respectively. Wild type data is shown in blue, while eRF1d data is shown in red, with significantly different bins highlighted in black. (d) Gene Ontology (GO) enrichment analysis was performed on all genes with significant ribosome occupancy changes at the −12 and −2 codon position, and GO terms related to translation (red) were significantly enriched, in addition to other terms (black). Analysis was performed with PANTHER, using Fisher’s Exact test, with a false discovery rate (FDR) correction. (e) In addition to translation termination, the process of cytoplasmic translation initiation (GO:0001732) was highly enriched, with nine out of a possible ten genes in yeast represented, including key members of the eIF2, eIF3, and eIF5 complexes. (f) Genes associated with both initiation and elongation (GO:0006414) were also highly enriched (15 out of 30).

RiboDiPA can not only identify the differential patterns of ribosome binding, but also important genes for downstream analysis. When we analyzed the identity of the approximately 850 genes that show DP at the −12 or −2 codon position (stop codon defined as −1 position), we found many interesting genes that are involved with the process of translation itself. Gene ontology (GO) term enrichment analysis showed significant enrichment for terms involved in protein translation among others (Figure 5d-e, *q*-value≤ 0.05). As expected, genes associated with translation termination were highly enriched, but also cytoplasmic translation initiation, including multiple members of the eIF3 and eIF5 complexes, and translation elongation, including members of the eEF1 family. Of the ten genes in yeast associated with cytoplasmic translation initiation, nine were pulled out in this analysis (Figure 5e), while half of genes associated with translation elongation were also identified. Taken together, these results demonstrate that DP analysis at single-codon resolution can discover differential patterns with fine details at the codon level, individually or collectively, as well as uncover putative regulatory feedback relationships in translation (see Section: DISCUSSION).

### 3.5 DP analysis for higher organisms

RiboDiPA can be readily applied to any other genomes including mammalian genomes. To this end, we examined data from [23] in mouse embryonic stem cells, and were able to discover many genes that showed differential patterns when comparing Nat1 heterozygous cells to Nat1 null cells. Figure S.2 in the Supplementary Materials shows two significant examples for Cdc20 and Gtf2f1, which are key regulators involved in cell cycle control and transcription respectively, and are both multi-exonic genes. Lower RPF coverage can be an issue for a single-codon resolution analysis for large genomes as too many bins may have extremely low counts. Therefore we recommend to use single-codon DP analysis only when coverage permits, and use the default adaptive binning when coverage is sparser, as is frequently the case in ribosome profiling experiments in mammalian systems.

### 3.6 Simulation Study

To further assess the performance of the proposed approach, we carried out a simulation study as follows. We constructed a two-condition Ribo-seq comparison experiment with *m* = 2, 3, 4 replicates within each condition respectively. To mimic the gene length and read count distributions observed in the real data, we selected the top 4000 genes that had largest total read counts under comparison of WT yeast cells versus eRF1 deletion strain in yeast data as templates. We randomly chose 3,600 genes as the null condition and the rest 400 as alternatives. For a given selected gene designated as the template for the null gene, we generated ribosome footprint data at the codon level by simulating the read count for each codon within each replicate using a negative binomial model *NB*(*µ*_*ijk*_, *ρ*_*ijk*_) where *i, j, k* are indices for replicate, codon and condition respectively. To do that, we first calculated the relative mean *q*_*j*_ for the *j*th codon based on all replicates. To generate the gene- and replicate-specific abundance/sequencing depth normalizing constant *s*_*ik*_ defined in (3) in a *m* vs. *m* comparison (i.e., *i* = 1, …, *m, k* = 1, 2), we first generated 2*m* random values from uniform[1,5] for each gene separately. Denote these random values as 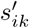. The normalizing constant *s*_*ik*_ is given by 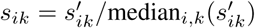. In this way, we allow a possible range of abundance fluctuation between 0.2 and 5 for each gene. The mean parameter is given by *µ*_*ijk*_ = *s*_*ik*_*q*_*j*_. In the second step, we generate the dispersion parameter *ρ*_*ijk*_ by plugging *q*_*j*_ into the fitted dispersion-mean curve function obtained from the wild type vs. eRF1 deletion comparison using DESeq2 package (i.e., output from *“estimateDispersionsFit”* R function within DESeq2).

For the 400 alternative genes, we randomly chosen 200 genes to contain 10% of codons with differential patterns, and the other 200 genes to bear 20% of codons with differential patterns. For a given selected template gene, we treated *q*_*j*_ for the *j*th codon of all replicates as the relative mean for the first condition, i.e., *q*_*j*1_. Considering that differential codons tend to be clustered in real data, we applied a Markov Chain model to simulate a sequence of the no-change or up-/down-regulated state path (see Supplementary Methods). If the codon is chosen to have a differential pattern with up-regulation, the relative mean for condition 2 is given by *q*_*j*2_ = *cq*_*j*1_, where *c* = 2, 4, 8 (or c=1/2, 1/4, 1/8 for down-regulation) in three separate simulation settings. If a codon is chosen to have no differential pattern, then *q*_*j*2_ = *q*_*j*1_. The dispersion parameter was generated in the same way as for the null genes.

In summary, our simulation is a 3 by 3 design, three different samples size (*m*) by three different log_2_-fold change with 10% true positive genes. We investigate how sample size, effect size and scale of differential patterns may affect the performance of the proposed approach.

We performed differential pattern analysis on both adaptively binned and unbinned data. We first examine the power achieved at nominal FDR level 0.05 of each setting (Figure 6). For both binned and un-binned data, the power consistently improves as sample size (*m*) or log_2_-fold change increases, while the DP analysis based on binned data achieves significantly better power than the codon-level analysis in each setting (Figure 6a). For example, the codon-level analysis achieved 47% power when the mean difference was 4 fold (lfc = 2) with 2 replicates and 88% with 4 replicates, whereas the binned analysis achieved 70% and 92% power respectively at the same settings. While at lfc=1 and m=2, only 5% and 15% power were achieved in the un-binned and binned analyses, suggesting small difference is difficult to decouple from the larger dispersion at small sample size and typical sequencing depth in current Ribo-seq studies.

**Figure 6:**
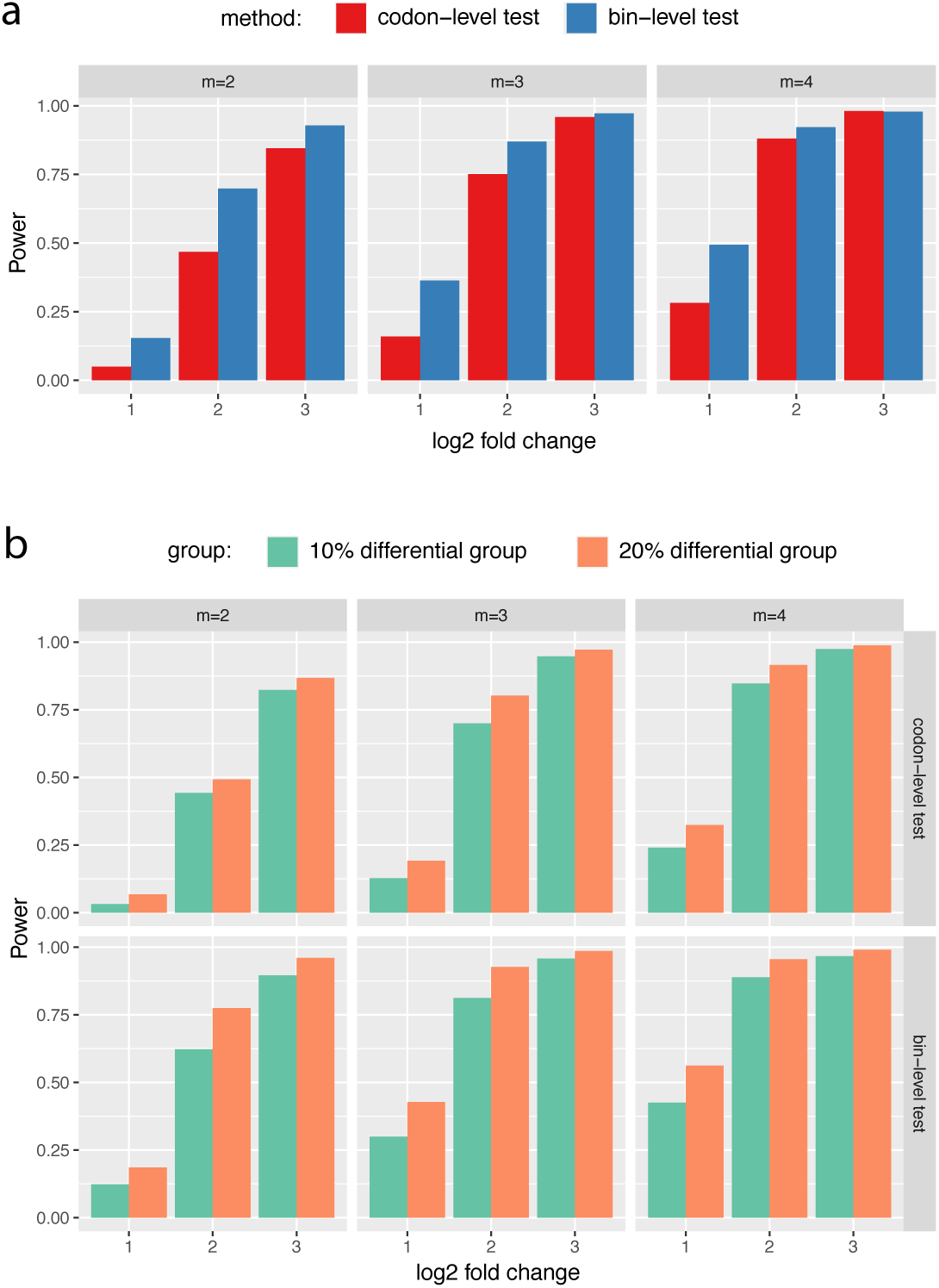
Simulation results I: (a) Power comparisons between binned and unbinned data (codon-level) in the simulation at nominal FDR level 0.05. (b) Power comparisons between groups of genes with 10% and 20% differential codons. The log2 fold change (lfc) of relative means between conditions of the true positive set was varied from 1 to 3, the number of biological replicates (*m*) was varied from 2 to 4. The presented results were averaged over ten repeated simulations.

We further split the power curve for the alternative genes with 10% and 20% differential codons (Figure 6b). Unsurprisingly genes with 20% of codons with differential patterns achieved larger power to be detected at the given sample size at nominal FDR level 0.05. Note the gene-level *p*-value from the multiple comparison procedure is determined by the minimum adjusted codon-level *p*-value across all codons within the same gene. In each simulation setting, regardless that the log_2_-fold change of *q* was fixed, more differential codons tend to result in more extreme minimum adjusted codon-level *p*-value and thus smaller gene-level *p*-value and larger power.

This simulation study provides new insights on the use of *T*-value as a supplementary measure for pattern differences beyond statistical significance. A larger pattern difference could be defined as a larger change of the relative mean *q*_*ijk*_ at one given codon/bin, or it can refer to more codons/bins/regions that had differential ribosome occupancy. The gene-level *p*-value is a sensitive measure for the first situation while *T*-value is informative for the latter. We compared ECDF or gene-level *p*-value and the *T*-value of the true alternative genes with 10% and 20% differential bins in all 9 settings in our simulation (Figure S.3). At lfc = 1, ECDF curve of *p*-value from the 20% group shows a leap over the 10% group at the left lower end (Figure S.3a,d,g), same for the T-value ECDF at larger *T*-value end (Figure S.3j,m,p). While the distinction of *p*-value curves diminishes at lfc=2 and 3 (Figure S.3b,e,h), the *T*-values of the 20% group maintain a significant gap over the 10% group. This demonstrates that *T*-value can be used as a supplementary measure to identify genes that have differential patterns over relatively larger regions.

We also compared Freedman-Diaconis rule with other binning methods including Sturges’ formula, Doane’s formula, Rice Rule, Scott’s normal reference rule, and an improved Doane’s formula. The ROC curve plots show that Freedman-Diaconis rule consistently outperforms all other methods under consideration in all simulation settings (Figure S.4).

## 4 Discussion

In this paper, we have proposed a novel statistical framework for differential pattern analysis for Ribo-seq data. DP analysis was defined for testing parallel mean curve between different conditions. This test allows for quick identification of genes that have differential ribosome binding patterns with rigorous quantification of statistical significance. We showed that DP analysis can effectively uncover global differences of translation between conditions, as well as discover fine-detail pattern differences up to single-codon resolution in genes that may play important roles in translational regulation.

One particular challenge in DP analysis in Ribo-seq data arises due to small sample size and low read counts at individual codons.. A typical Ribo-seq experiment often contain 1 or 2 replicates per condition. In our simulation, we showed that with two replicates and 2 fold change of mean pattern difference, we only achieved 5% and 15% statistical power (sensitivity) for unbinned (codon-level) and binned data when controlling the nominal FDR ≤ 0.05 (Figure 6a). There are multiple factors that may contribute to the poor power. First, the small effect size is difficult to decouple from the large variance when the sample size is small. Second, the fitted dispersion-mean curve typically has larger dispersion for small means, which may further exacerbate the power for genes with small read counts (even after binning). We defined the differential pattern in terms of the relative mean at every individual codon/bin in the entire coding region. An advantage of this framework is that it allows one to pinpoint differential patterns at any specific bin/location. A downside is that the codon-wise/bin-wise read count can be too low to detect true difference. Furthermore the point-wise testing procedure requires a family-wise type I error correction for gene-level *p*-value calculation. The hybrid Hochberg-Hommel procedure improves over Bonferroni and some other correction methods in power, but still tends to be conservative overall.

Ribo-DiPA provides a new angle to investigate translational regulation using Ribo-seq data. Unlike most of existing methods that use total read counts to quantify translation efficiency, Ribo-DiPA examines the actual ribosome footprint distribution along the transcript, which may provide additional insights into the fine-scale translational activities. When we compared data from WT yeast cells with cells where the translation terminator eRF1 was removed, approximately 600 genes showed significant differences at −12 and −2 codons upstream of the stop codon (see Figure 5a). The ternary complex formed by eRF1 with eRF3 and GTP which is plays an essential role for translation termination in eukaryotes, and in yeast, eRF1 recognizes all three canonical stop codons [36]. Since the vast majority of transcripts require this complex in order to terminate translation, *a priori* it was possible that the genes with DP in the eRF1d condition would reflect the most highly translated genes in yeast, since higher translation would result in more ribosome complexes that are unable to finish translation. Instead we found many genes with significant DP were not the most highly expressed genes in yeast (i.e. top 600 genes by RPF abundance in wild type), although there was some overlap. This suggests that the identity of this subset of genes is not merely a reflection of high expression.

Further, we noted a very striking enrichment of genes involved in cytoplasmic translation initiation in this condition, as well as translation termination and elongation (Figure 5d-f). We speculate that arresting ribosomes on these genes could serve as a mechanism to quickly reinitiate translation once the stalling stress has passed. In eukaryotes, stalled ribosomes are recognized by the Ribosome-associated Quality Control (RQC) pathway and quickly targeted for subunit dissociation, mRNA degradation, and nascent polypeptide degradation [37]. The recognition of stalled ribosomes is thought to be primarily mediated by ZNF598 (Hel2 in yeast), and involve recruitment of numerous downstream factors [38]. We speculate that in most cases, the formation of many stalled ribosomes as visualized in Figure 5a would lead to triggering the RQC pathway and subsequent degradation, but that ribosomal proteins and translation initiation factors might be protected from degradation and therefore are enriched in DP analysis under these conditions by being retained. This would allow for rapid reestablishment of translation after a global stalling stress had passed, although we cannot rule out that eRF1 depletion is an extreme perturbation that overwhelms the capacity of the RQC pathway. A recent paper demonstrated a link between the RQC pathway and translation initiation by showing ZNF598 collaborates with GIGYF2 and 4EHP proteins to inhibit translation eIF4G-dependent initiation of stalled messages [39]. By extension, we postulate that mRNAs for components of the translation machinery may be preserved from RQC in order to restart initiation, although the mechanism as to how these messages would forestall recognition by ZNF598 and degradation remains to be uncovered.

## 5 ACKNOWLEDGEMENTS

This research is supported by the NSF-Simons Center for Quantitative Biology (Simons Foundation/SFARI 597491-RWC and the National Science Foundation 1764421). The authors also want to thank Dr. Jiangtao Gou for helpful discussion on multiple comparison method.

