## supplementary materials for "RiboDiPA: A Novel tool for differential pattern analysis in Ribo-seq data"

#### S.1 Data acquisition

Data were downloaded from [1] and [2]. The table below lists the comparisons made throughout the paper, with corresponding GEO and SRA numbers.

**Yeast data:**

**WT yeast vs. eRF1d yeast (unstressed**

WT_CHX1 (GSM3168380; SRR7241903)

WT_CHX2 (GSM3168381; SRR7241904)

eRF1_depletion CHX replicate 1 (GSM3168385; SRR7241908)

eRF1_depletion CHX replicate 2 (GSM3168386; SRR7241909)

**Unstressed vs. Osmotic**

Wild-type CHXTIG profiling 1 (GSM3168396; SRR7241919)

Wild-type CHXTIG profiling 2 (GSM3168397; SRR7241920)

Wild-type hyperosmotic stress profiling 1 (GSM3168407; SRR7241930)

Wild-type hyperosmotic stress profiling 2 (GSM3168408; SRR7241931)

**Unstressed vs. Oxidative**

Wild-type CHXTIG profiling 1 (GSM3168396; SRR7241919)

Wild-type CHXTIG profiling 2 (GSM3168397; SRR7241920)

Wild-type oxidative stress profiling 1 (GSM3168403; SRR7241926)

Wild-type oxidative stress profiling 2 (GSM3168404; SRR7241927)

**Unstressed vs. Stationary Phase**

Wild-type CHXTIG profiling 1 (GSM3168396; SRR7241919)

Wild-type CHXTIG profiling 2 (GSM3168397; SRR7241920)

Wild-type stationary phase profiling 1 (GSM3168401; SRR7241924)

Wild-type stationary phase profiling 2 (GSM3168402; SRR7241925)

**WT Osmotic Stress vs. Rck2 Osmotic Stress**

Wild-type hyperosmotic stress profiling 1 (GSM3168407; SRR7241930)

Wild-type hyperosmotic stress profiling 2 (GSM3168408; SRR7241931)

rck2 hyperosmotic stress profiling 1 (GSM3168409; SRR7241932)

rck2 hyperosmotic stress profiling 2 (GSM3168410; SRR7241933)

**WT Oxidative Stress vs. Rck2 Oxidative Stress**

Wild-type oxidative stress profiling 1 (GSM3168403; SRR7241926)

Wild-type oxidative stress profiling 2 (GSM3168404; SRR7241927)

rck2 oxidative stress profiling 1 (GSM3168405; SRR7241928)

rck2 oxidative stress profiling 2 (GSM3168406; SRR7241929)

**Mouse embryonic stem cell data:**

**Nat 1 Het. vs Nat 1 Null**

Nat1+/- A7 RPF rep1 (GSM2357216; SRR4436402)

Nat1+/- B4 RPF rep2 (GSM2357220; SRR4436406)

Nat1+/- C1 RPF rep3 (GSM2357224; SRR4436410)

Nat1-/- B12 RPF rep1 (GSM2357222; SRR4436408)

Nat1-/- B2 RPF rep2 (GSM2357218; SRR4436404)

Nat1-/- C3 RPF rep3 (GSM2357226; SRR4436412)

#### S.2 Data pre-processing and alignment

Data were download using the SRAtoolkit using the `fastqdump --gzip` command. Cutadapt, ver1.14, was used to trim 3' adaptors from reads, with all data showing $> 90\%$ of reads with adaptors. For eRF1 deletion data [1] and its wild type comparison, the command : `cutadapt -q 10 -a CTGTAGGCACCATCAA` were used. For yeast stress data [1], `cutadapt -u 4 -a NNNNNNCACTCGGGCACCAAGGA` were used to remove the 4 nucleotide 5' unique molecular identifier (UMI) and 3' 6 nucleotide UMI. For the Sugiyama data [2], the following command and parameters were used: `cutadapt -a AGATCGGAAGAGCACACGTCT`.

For depletion of rRNA sequence, bowtie2, ver2.2.6, was used to generate a custom genome of just rRNA sequences for alignment, and then unaligned reads were preserved for alignment to the genome. For the yeast genome, the command `bowtie2-build YeastRibosomeRNA.fasta Sc-rRNA` was used, while for the mouse genome the command `bowtie2-build Mm-rRNA-combined.fasta Mm-rRNA` was used. For yeast rRNA, a pre-rRNA sequence containing 5.8S, 18S, and 28S rRNA sequences (RDN37-1, 35S pre-rRNA), as well as mature 5S rRNA was used. For mouse rRNA, only mature 5S, 5.8S, 18S, 28S, 12S, and 16S rRNA sequences were used. Ribosomal RNA files for both species were sourced from NCBI, and are available on request. Reads were aligned to the rRNA sequences using the command

```
bowtie2 --local -x ./ribo-index/Sc-rRNA \
--un-gz ./ribo-deplete/SRRXXXXXXX_mRNA.fastq.gz \
-U ./fastq-trimmed/SRRXXXXXXX_trimmed.fastq.gz \
-S ./ribo-deplete/SRRXXXXXXX_rRNA.sam
```

and

```
bowtie2 --local -x ./ribo-index/Mm-rRNA \
--un-gz ./ribo-deplete/SRRXXXXXXX_mRNA.fastq.gz \
-U ./fastq-trimmed/SRRXXXXXXX_trimmed.fastq.gz \
-S ./ribo-deplete/SRRXXXXXXX_rRNA.sam
```

for yeast and mouse respectively. The files `SRRXXXXXXX_mRNA.fastq.gz` were taken for further alignment.

Suffix array genome files were generated using STAR, ver2.6.0, with yeast reference genome assembly SacCer3, and mouse genome assembly Mm10. GTF and FASTA files were sourced from ENSEMBL:

```
STAR --runMode genomeGenerate --runThreadN 12 --genomeDir . --genomeFastaFiles \
../genome-files/Saccharomyces_cerevisiae.R64-1-1.dna.toplevel.fa --sjdbGTFfile \
../genome-files/Saccharomyces_cerevisiae.R64-1-1.98.gtf --sjdbOverhang 49 \
--limitGenomeGenerateRAM 100000000000
```

and

```
STAR --runMode genomeGenerate --runThreadN 12 --genomeDir . --genomeFastaFiles \
../genome-files/Mus_musculus.GRCm38.dna.toplevel.fa --sjdbGTFfile \
```

```
../genome-files/Mus_musculus.GRCm38.96.gtf --sjdbOverhang 39 \
--limitGenomeGenerateRAM 100000000000
```

Data were aligned with STAR, using the flags "–quantMode GeneCounts –outFilterScoreMinOverLread 0.3 –outFilterMatchNminOverLread 0.3 –outFilterMultimapNmax 100". For eRF1d and companion comparison data, read length was filtered to only include reads 20 to 32 nucleotides in length.

#### S.3 P-site mapping

We followed [1] to infer the P-site location based on the the following empirical rules: 20:[13], 21:[14],22:[14], 27:[13], 28:[13], 29:[14], 30:[14], 31:[14], 32:[14], where the number before ":" is the RPF length and number inside of brackets is the P-site offset inferred from the 5' end of RPF.

#### S.4 Simulation of differential pattern

In real data, codons with differential patterns tend to be clustered. Within the same cluster differential codons tend to have the same up-/down-regulated pattern. For true alternative genes, each codon can take one of the three states: no differential pattern (N), up-regulated pattern (U) and down-regulated pattern (D). To simulate the sequential dependence of the three states for each gene that contains differential pattern, we considered a Markov chain model with transition probabilities specified as follows:

$$P_1 = \begin{bmatrix} 0.96 & 0.02 & 0.02 \\ 0.35 & 0.65 & 0 \\ 0.35 & 0 & 0.65 \end{bmatrix}, \quad P_2 = \begin{bmatrix} 0.910 & 0.45 & 0.45 \\ 0.35 & 0.65 & 0 \\ 0.35 & 0 & 0.65 \end{bmatrix},$$

where the stationary distributions of $P_1$ and $P_2$ are roughly $(0.9, 0.05, 0.05)$ and $(0.8, 0.1, 0.1)$ respectively. The initial probabilities for "N", "U", "D", are set as $p_1 = (0.9, 0.05, 0.05)$ and $p_2 = (0.8, 0.1, 0.1)$ for genes with $10\%$ and $20\%$ differential bins respectively. As such we will control the total differential codons at $10\%$ and $20\%$ while imposing positive correlation of differential pattern status among nearby codons.

#### S.5 Data binning methods comparison

Suppose there are $m$ P-sites mapped at locations $x_1, \ldots, x_m$ of a transcript. We compared the following six adaptive binning approaches ($k$ is the bin number and $h$ is the bin width below):

- Sturges' formula [3]: $k = \log_2 m + 1$, where $k$ is the choice of number of bins.
- Doane's formula [4]: $k = 1 + log_2 m + log_2(1 + |g_1|/\sigma_1)$, where $g_1$ is the 3rd moment skewness, and $\sigma_1 = \sqrt{(6(m-2)/((m+1)(m+3))}$.
- Rice Rule [5]: $k = 2m^{1/3}$.
- Scott's normal reference rule [6]: $h = 3.49sd/m^{1/3}$, where $h$ is the choice of bin width and $sd$ is the standard deviation.
- Freedman–Diaconis' choice [7]: $h = 2IQR/m^{1/3}$, with IQR is the interquartile range of the data. It is robust against outliers in data.

- An variant of Doane's formula using a kurtosis criterion [8, 9]: $k = \min\{\log_2(m) + 1 + \log_2(1 + K \cdot \sqrt{m/6}), m\}$, with $K = \sum_{i=1}^{m}(x_i - \bar{x})^4/[(m-1)sd^4]$.

### S.6 Supplementary figures and legends

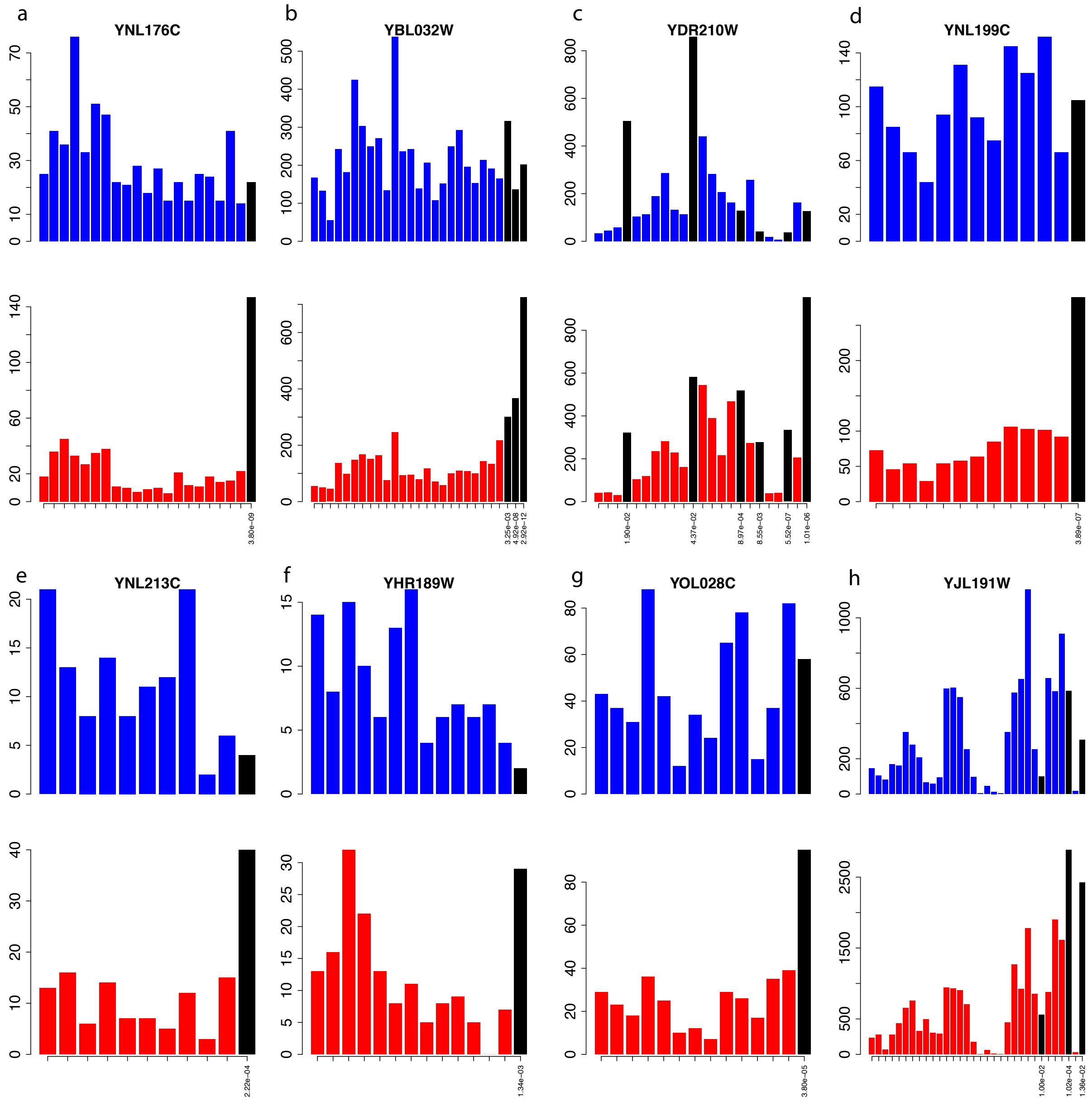


Figure S.1: **DP analysis for WT vs. eRF1 deletion strain for yeast data with adaptive bin width**. (a)-(h) Eight genes were selected with one replicate for each condition to show similar up-regulation pattern of P-sites near stop codon in eRF1. WT and eRF1d conditions are colored blue and red data respectively. Bins with significant differences in pattern between conditions are colored black.

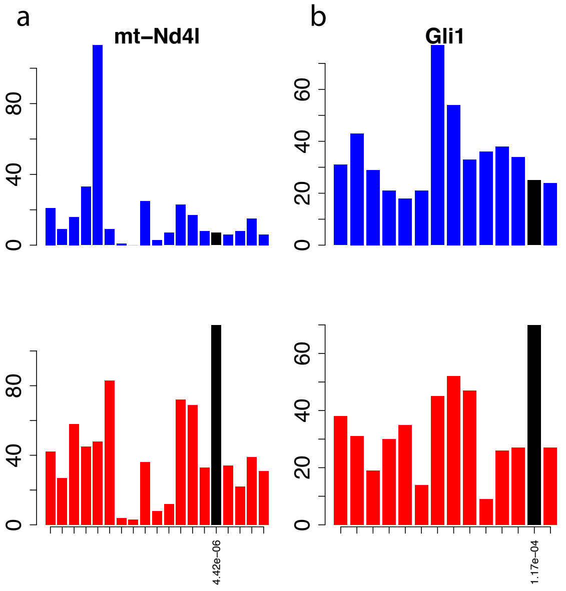


Figure S.2: **RiboDiPA DP analysis for mammalian organism**. Data in this figure were adapted from [2], comparing mouse embryonic stem cells, with various genetic dosages of the translational regulator Nat1 heterozygous mutant (blue), versus null (red). Coloring are the same in Figure S.1. (a) mt-Nd4l (ENSMUSG00000065947), and (b) Gli1 (ENSMUSG00000025407).

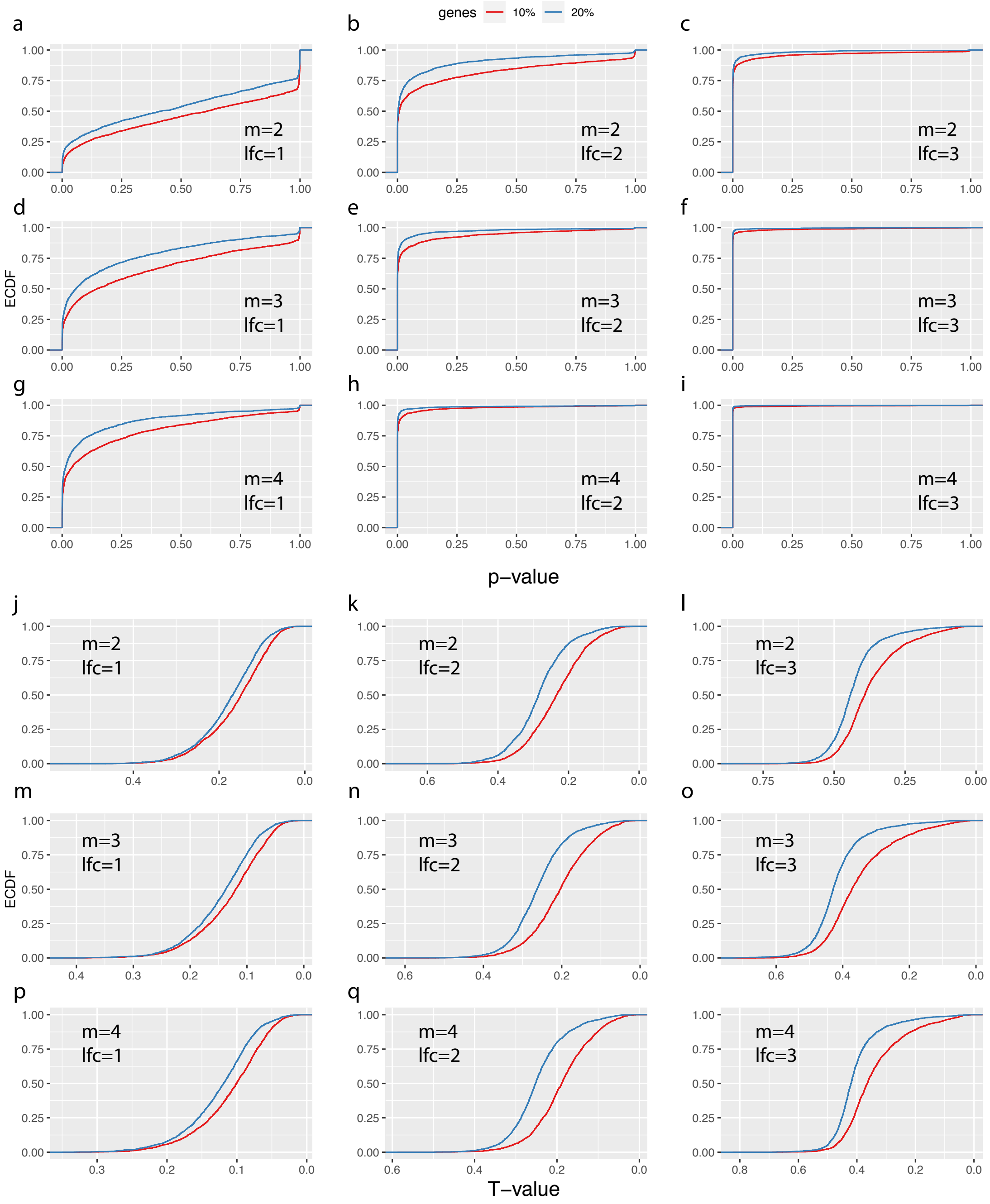


Figure S.3: **Simulation results II: $T$-value is effective metric to identify genes with larger pattern difference**. (a)-(i) ECDF of $p$-value for true alternative genes in each simulation setting. Genes with 10% and 20% differential bins were plotted separately. (j)-(r) Same ECDF as in (a)-(i) but for $T$-value, plotted in descending order of $T$-value. $T$-value ECDF plot shows better distinction between the 10% and 20% groups demonstrating its effectiveness for identifying genes with larger DP regions. The presented results were averaged over ten repeated simulations.

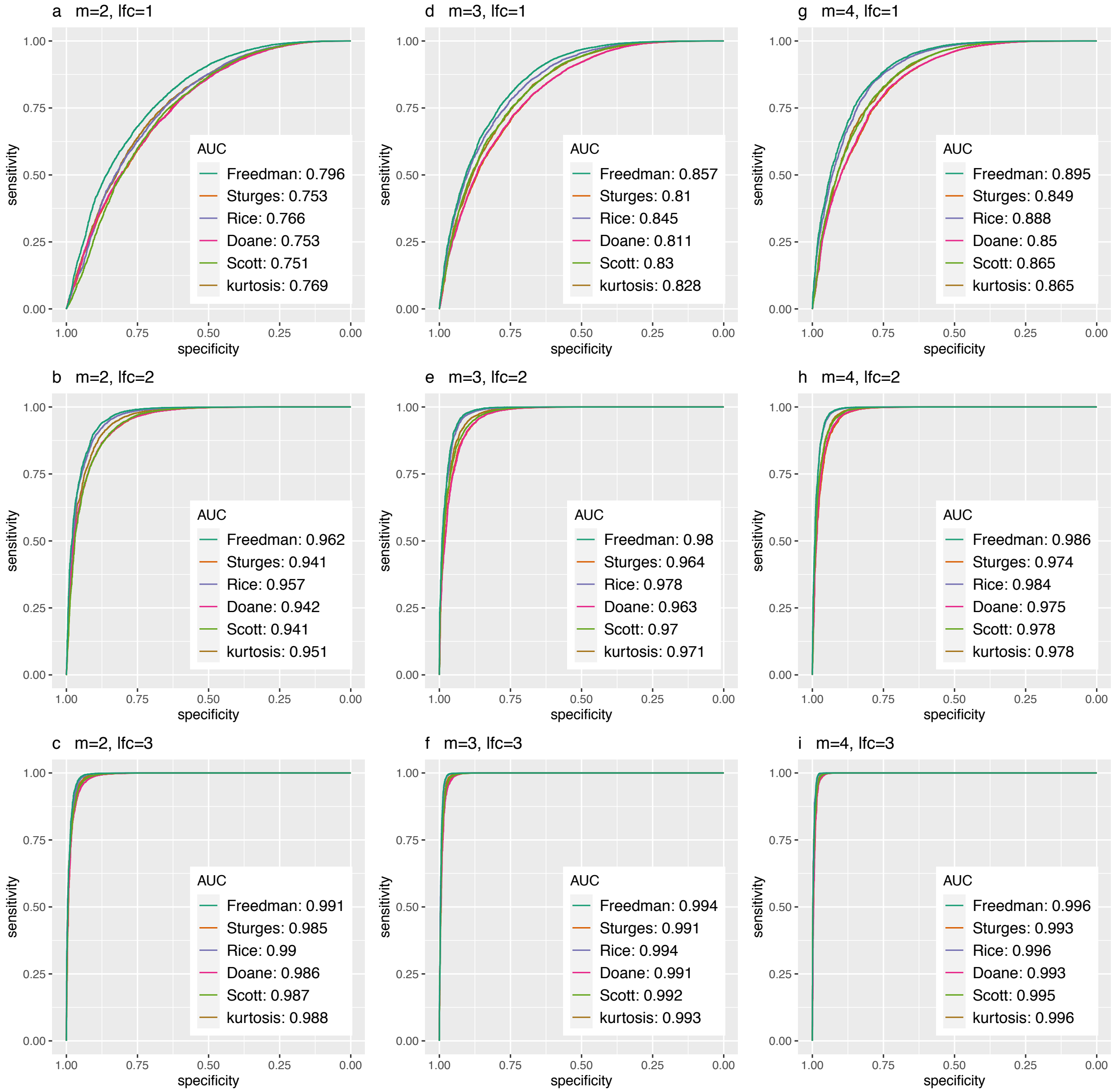


Figure S.4: **Simulation results III: Receiver operating characteristic (ROC) curves of different binning methods for all settings**. The log2 fold change (lfc) of relative means between conditions of the true positive set was varied from 1 to 3, the number of biological replicates ($m$) was varied from 2 to 4. Results are averaged over ten repeated simulations.
